# Nudibranch color diversity shares a common origin in guanine photonic structures

**DOI:** 10.1101/2025.09.06.674434

**Authors:** Samuel Humphrey, Xianglian He, Tobias Priemel, Vera Titze, Sinuhé Perea-Puente, Vivek Subramanian, Cedric Bouchet-Marquis, Bruno Jesus, Silvia Vignolini

## Abstract

Nudibranchs are well known for their bright and diverse color patterns. This coloration is typically a form of aposematism, warning predators against toxic compounds sequestered from their prey and weaponized as a form of defense. Although many of the hues in nudibranchs have pigmentary origin, multilayer structures composed of guanine nano-platelets have been suggested as the source of color enhancement in the nudibranch *Flabellina iodenea*. Here, using a combination of white light and Raman microspectroscopy techniques, we report that such guanine-based multilayer structures are a widespread mechanism to create angular-independent structural color across the dorid and aeolid groups. Additionally, by using cryo-FIB tomography, we were able to access the complex 3D organization of the guanine nano-platelets responsible for the strong blue coloration of *Chromodoris annae*. We propose that the multilayer organization of guanine platelets with varying orientations across the tissue offers a particularly effective strategy for producing diverse optical effects. In this configuration, hue is mainly governed by interlayer spacing, while the angular dependence of color can be tuned through the degree of local order, allowing a single structural motif to generate a broad palette of optical appearances.

**Significance statement:** Nudibranchs are an extraordinarily diverse group of marine animals, renowned for their dazzling range of colors and striking patterns. Whilst their pigmentary coloration is well understood, so far, structural coloration, obtained only by nanostructures, has only been reported in the nudibranch *Flabellina iodenea*. In this work, we present a comparative analysis of structural coloration across nudibranch species from both benthic and coral reef environments, and we show that guanine-based nano-structures are a common motif responsible for a wide range of colors, spanning the dorid and aeolid groups. We foresee that the 3D imaging conducted here may serve as inspiration for bio-photonics studies in other marine organisms, and that the structures themselves could serve as inspiration for bio-inspired materials.

## Introduction

Nudibranchs (Heterobranchia) exhibit some of the most brilliantly diverse colors and patterns of all marine organisms(1–3). This coloration is thought to have evolved with shell loss, in many cases as a form of aposematism, or camouflage(3, 4, 5). They are known to be prolific kleptochemists, stealing toxins, stings, and spikes from their prey organisms. Notable examples include the ‘blue dragon’ sea slug *Glaucus atlanticus*, which steals stinging cells (nematocysts) from the Portuguese Man o’ War Jellyfish, transferring them to its own tissue to use as a defense mechanism(6), and the ‘Spanish dancer’ nudibranch *Hexabranchus sanguineus*, which steals chemical defenses from its sponge prey(7). A strong correlation has been shown to exist between conspicuousness and toxicity in nudibranchs(4, 8), and 50% of opisthobranchs are believed to show aposematic coloring(8).

Whilst the extraordinary variety of their color palette is well documented, and several pigments have been proven responsible for such hues(9, 10), coloration coming from nanostructures, so-called structural color, has been largely overlooked in these species. So far, structural color has only been reported in one species of aeolid nudibranch, *Flabellina iodenea* (5). In this species(5), guanine crystal stacks were observed in the tips of cerata and rhinophores, and such structures are suggested to be responsible for enhancing the vividness of the pigmentary color pattern through ‘silvery’ reflectance, as well as increasing visibility of the orange cerata at depths where penetration of light at the red end of the spectrum is low. When observed in situ through TEM, envelopes near the epithelial basal lamina contained guanine platelets free within cells and membrane-bound. These platelets vary in size between 200-800 nm across the flat face of the crystal, and 50 – 200 nm in the stacking direction, in stacks of 2 to 8.

Interestingly, guanine-based structural color is found in a wide range of organisms from vertebrates, such as panther chameleons(11) and fish (12–15), to invertebrates such as spiders(14) and mollusks (e.g. the image-forming mirrors in the eyes of scallops(16)). The crystal structure of β-guanine (the form typically found in nature) can be seen in **Supplementary Figure 1**.

Here, we have studied a range of dorid and aeolid nudibranchs from across their phylogenetic tree, showing that guanine photonic structures are a common thread to produce structural coloration. These species were chosen based on their optical appearance and diversity within the two selected groups. Dorid and aeolid nudibranchs exhibit a wide range of colors and patterns, but with some key similarities (see **Supplementary Figure 2**). The coloration in these groups is often pixelated (composed of discrete granules), described as ‘vivid’, ‘silvery’ or ‘iridescent’, and encompasses colors which, in other animals, are typical of structural coloration (white, violet, blue)(1, 17, 18). Additionally, pigmentary analysis has thus far only been able to account for specific hues in species within patterns of multiple colors, suggesting pigmentary color may not be the only mechanism(9, 10). Other groups, such as Dendronotoidea and Arminida, have not been selected as they do not display such features and are typically dull in color. The range of colors with similar optical properties, but varied patterns and hues across the dorid and aeolid groups raises the question of whether the same color generation mechanism is utilized in both cases.

To characterize the optical response and composition, and to understand the role of guanine crystals in the optical appearance of nudibranchs, we used white-light micro-spectroscopy and Raman spectroscopy on histological sections, respectively. Finally, we revealed the full 3D hierarchical nano-architecture responsible for *Chromodoris annae*’s structural coloration, using cryogenic focused ion beam scanning electron microscopy (cryo-FIB SEM). This imaging is the first of its kind for structural coloration, and we foresee that it can be useful for visualizing the nanoarchitecture of guanine in a wide range of living organisms due to guanine’s excellent contrast in the secondary electron mode. We believe that it is not by chance that guanine nanostructures with different degrees of order are responsible for the optical appearance of a group of animals known to be one of the most chromatically diverse on earth. Such nanostructures can provide a wide range of appearances, both in terms of hue and angular dependence by simply tuning the geometry of the multilayer structure in periodicity, size, and polycrystallinity. The striking coloration of nudibranchs plays a key role in predator deterrence and so the capability of photonic structures to generate distinct, bright hues may confer color diversity.

## Results and discussion

### Structural color variation in nudibranchs

We selected 14 nudibranch species (6 dorids, 8 aeolids) and constructed a phylogenetic tree based on NCBI taxonomic data (**Figure 1A**) to highlight the distribution of color traits across their evolutionary history (see methods section for details of construction)(19). From this broader set, we focused on six representative species — *Hypselodoris tryoni, Hypselodoris bullockii, Chromodoris annae, Chromodoris willani, Spurilla neapolitana, Berghia stephanieae* — chosen for their color diversity and their phylogenetic range within the dorid and aeolid clades. As visible in **Figure 1**, pixelated, iridescent colored granules are a common thread in these species, and so, despite their varied macroscopic appearance, we expected that the mechanism of color production may be the same. Additionally, all the above dorid nudibranchs sequester secondary metabolites as chemical defenses (20–22), whilst the aeolid nudibranchs utilize stolen cnidocytes from anemones (23–25), so structural color in this context may have an aposematic function in all of the species studied here.

**Figure 1.**
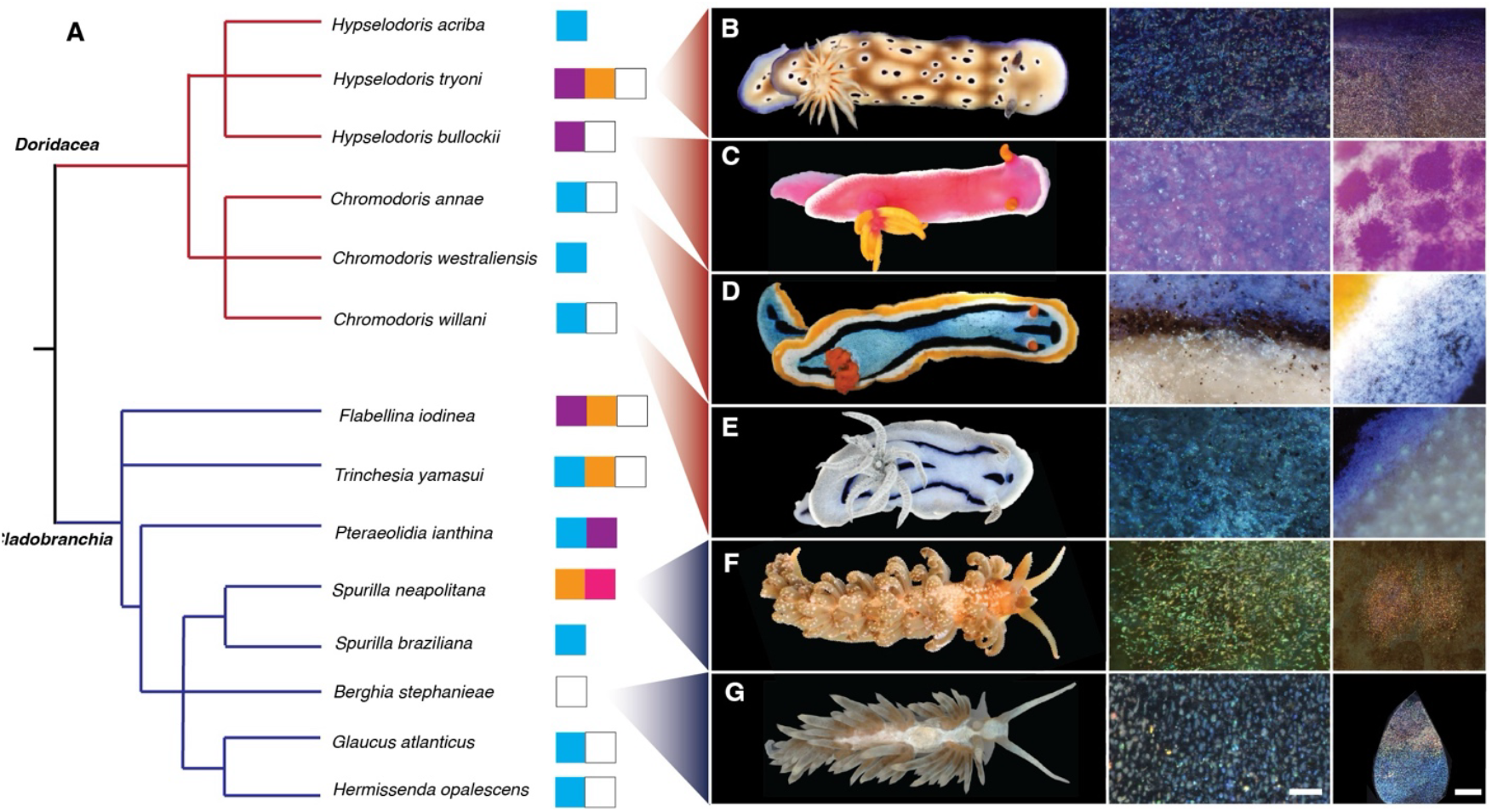
A) Phylogenetic tree containing 14 nudibranch species from the clades Doridacea and Cladobranchia. The species studied here are visible in the photographs on the right (from top to bottom [B-G]: *Hypselodoris tryoni, Hypselodoris bullockii, Chromodoris annae, Chromodoris willani, Spurilla neapolitana, Berghia stephanieae)*. Other species have either been reported to utilize guanine crystals or were selected based on their visual appearance (iridescent, brilliant, pixelated coloration on the skin) in order to show the diversity of genera believed to utilize guanine for structural color within the two highlighted clades. Images of these species can be found in **Supplementary Figure 2**. The phylogenetic tree was constructed using taxonomic data from the NCBI taxonomy database. Colored squares show the range of macroscopic colors believed to be the result of guanine crystal based structural color for each species (i.e. pigmentary colors are not included). B-G) Digital microscope images showing structurally colored granules in *Hypselodoris tryoni*, skirt (B), *Hypselodoris bullockii* mantle (C), *Chromodoris annae* skirt (D), *Chromodoris willani* mantle (E), *Spurilla neapolitana* ceras (F), *Berghia stephanieae* ceras (G). (Scale bars, 50 *μ*m, middle column; 200 *μ*m, right column.)

In the four dorid nudibranchs shown here, the skirt coloration is distinct from the mantle. In all cases, there are regions of white coloration and regions of violet or blue coloration. For *H. tryoni* the outer skirt is colored in a violet hue, whilst the mantle is white/beige colored with mauve spots, *H. bullockii* has a white skirt with pink/violet coloration on the mantle and at the base of the rhinophores and yellow gills/rhinophores, *C. annae* has 5 distinct regions of coloration with stripes of yellow, white, black and blue on the mantle and orange gills/rhinophores, *C. willani* has a white skirt and black/blue stripes on the mantle. The two aeolid nudibranchs shown here have coloration concentrated on their cerata, in the case of *S. neapolitana* in the form of orange spots, whilst for *B. stephanieae*, the entire cerata appear white (however, at high magnification can be seen to show a gradient of coloration (**Figure 1G**, right-hand column). *Spurilla neapolitana* is colored brown by *Symbiodinium* microalgae sequestered from its anemone prey.

To confirm the structural origin and characterize the optical properties of coloration in the above species, high magnification optical imaging in the Köhler illumination configuration was done on the structurally colored regions shown in **Figure 1**. These regions can be clearly recognized in the microscope images of **Figure 2A-H**. In all cases, the color in the images appears pixilated; composed of discrete regions of different color with diameters ranging from 0.2 to 16 μm,. Grain diameter varies between species: *H. tryoni –* 3 ± 2 μm, *H. bullockii –* 7 ± 4 μm, *C. annae –* 3 ± 3 μm, *C. willani –* 1.0 ± 0.7 μm, *S. neapolitana –* 5 ± 2 μm, *B. stephanieae –* 8 ± 4 μm. *C. annae* and *C. willani* display blue coloration together with black pigmentation to increase contrast(26). Such small granules reflecting all colors will behave like pixels, at a microscopic scale; therefore, due to color mixing, the final observed resolution will be the intensity-averaged sum of the color reflected from individual pixels. This effect is like the function of light-emitting diodes (LEDs) in an RGB display (27) and has also been observed in other species, such as the fruits of *Pollia condensata*(28).

**Figure 2.**
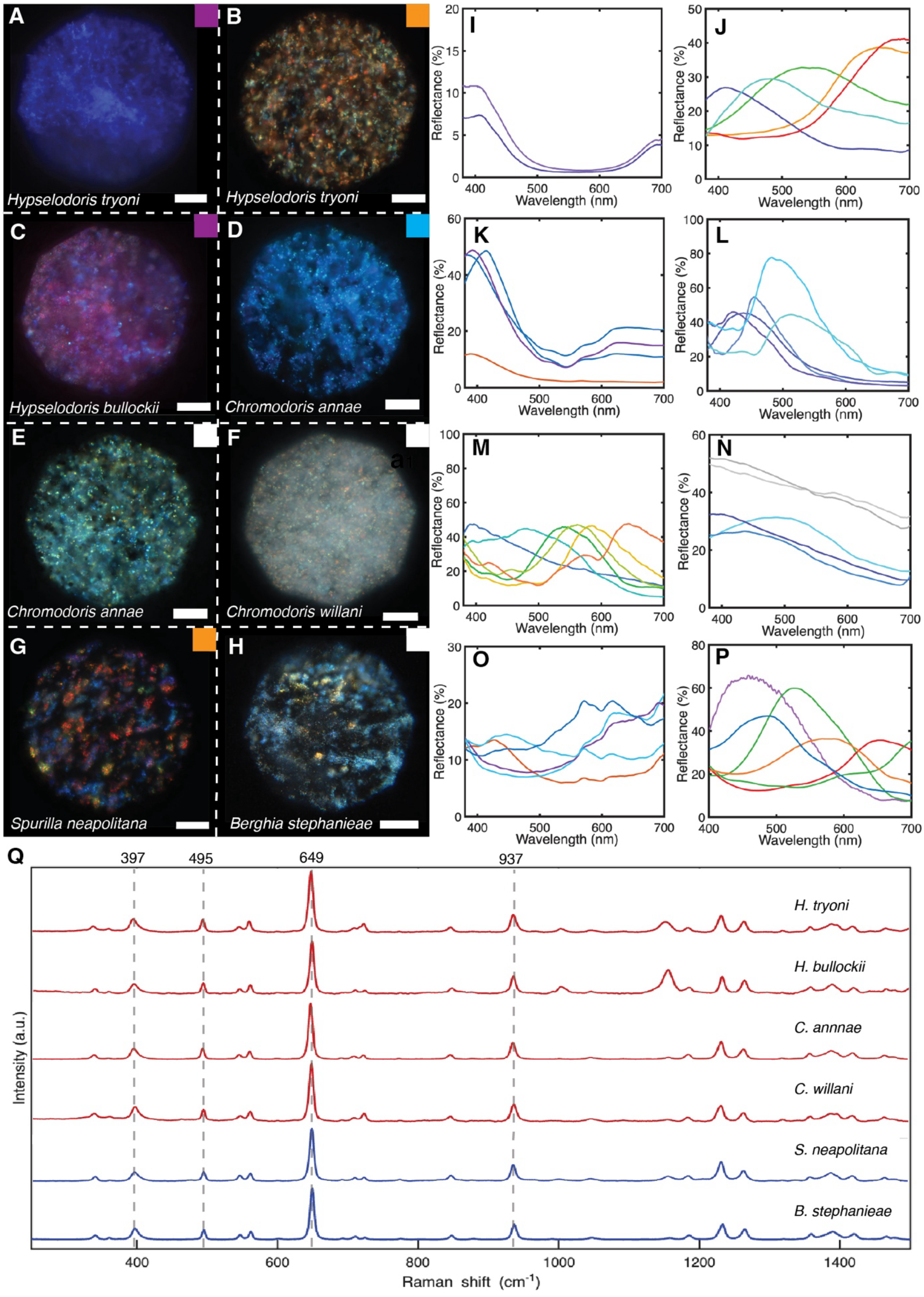
A-H) Brightfield upright optical microscope (Zeiss axioscope, 40x WI objective) images of structurally colored regions on the dorsal surface of the 6 species shown in Figure 1a. Scale bars, 30 *μ*m. The color of the squares at the left top corner of the image corresponds to the structural coloration perceived macroscopically. A) *Hypselodoris tryoni* violet region, located at the perimeter of the skirt. Only violet domains of consistent color are visible here. B) *Hypselodoris tryoni* beige/white region. Domains of all colors are visible here, with a higher frequency of those in the orange-red wavelength range. C) *Hypselodoris bullockii* pink region. Domains of violet and background pink coloration are visible. D) *Chromodoris annae* blue region showing granular appearance and cloudy patches. E) *Chromodoris annae* white region, showing multiple domains of all colors across the visible spectrum. F) *Chromodoris willani* white region. Colored domains across the visible spectrum are visible atop a cloudy white background. G) *Spurilla neapolitana* orange region (located on the cerata) showing discrete domains of color across the visible wavelength range. H) *Berghia stephanieae* ceras. Colored domains across the visible spectrum are visible, dominated by those at the blue end of the spectrum. In all of the above images, colored domains ranging in size from 0.2 - 16 *μ*m appear to have random orientation. I-P) Brightfield reflectance spectra (Zeiss axioscope, 40x WI objective, 50 *μ*m fibre) normalized to a silver mirror, corresponding to the colored regions shown in A-H. Peak shape, intensity, and position vary between species. Notably, white regions (*C. annae* [E], *C*. willani [F]) appear to have spectral peaks across the visible range, suggesting a coloration mechanism similar to an LED in which colored pixels mix to give an overall white appearance. Orange/beige regions (*H. tryoni* [B], *S. neapolitana* [G]) also show spectral peaks across the visible range, although dominated by those in the orange-red wavelength range. *C. willani* shows a broadband white spectrum when the focal plane is changed to the ‘cloudy’ material. Q) Raman spectra of iridescent regions located close to the dorsal surface in histological sections of the 6 nudibranch species studied here. Dashed lines indicate peak positions of literature values for anhydrous biogenic guanine.

Reflectance spectra of individual granules were measured (all spectra are referenced to a silver mirror) for the different species. As visible in **Figure 2A-H**, different species and different regions within one species show different reflections. For example, the blue region in *C. annae* (**Figure 2D**) has reflectance peaks with up to 80% intensity centered at wavelengths within 420 – 515 nm (**Figure 2L**), whilst for the white region (**Figure 2E**), the reflectance peaks span across the visible spectrum (395 – 645 nm) with a consistent reflectance of ∼ 50%, leading to a white macroscopic appearance. *Hypselodoris tryoni* again has two distinct structurally colored regions (**Figures 2A, B**), a violet one (reflectance peaks from 395 – 410 nm ([**Figure 2I)]**) and a white/orange region, where we observed reflectance peaks from 410 – 690 nm (**Figure 2J**), weighted towards those at longer wavelengths. This intensity difference will result in an orange/white appearance macroscopically, despite individual granules also reflecting shorter wavelengths. At high magnification, *H. bullockii* displays violet granules above a pink background (**Figure 2C**), which results from a layer of pigment lying below the granules. The violet granules possess reflection spectra with intensity up to 50% in the wavelength region from 385 – 415nm (**Figure 2K**). *Spurilla neapolitana* displays granules with color across the visible wavelength range (**Figure 2G**). Spectral peaks are weighted by intensity towards those at longer wavelengths (**Figure 2O**) to give an orange macroscopic appearance. The white region in *C. willani* is composed of two layers, which can be imaged at different focal planes (**Supplementary Figure 3**). In **Figure 2F** we see structurally colored granules with reflectance across the visible range (corresponding to the blue lines in **Figure 2N**) whilst in the focal plane lying above these granules, there is a diffuse white material (corresponding to the grey spectra in **Figure 2N**). We hypothesize that the diffuse material acts to scatter light reflected from the colored domains, homogenizing the colored reflections, to give a white, rather than silvery appearance, analogous to diffuse reflectors observed in butterfly wing scales(29). Whilst the coloration shown here is consistently composed of discrete, angular granules with random orientation (0.2 – 16 *μ*m diameter), the visual appearance of the animals varies widely. We propose that this occurs by the variation in spectral properties of the individual granules (due to differences in ultrastructure), and by the statistical distribution of these spectra when averaged over many granules.

To confirm the presence of biogenic guanine crystals as reported in the nudibranch *Flabellina iodenea*, we used Raman spectroscopy on histological sections of tissue from all the described nudibranchs: four dorid, two aeolid species. Raman is a convenient method to identify the material present in the section. Raman spectra were taken of areas that appeared to be highly reflective in the brightfield mode. Although the color of the 6 species is highly varied, the material used to produce this color is the same. The six species sampled have peaks at Raman shifts of 397, 495, 649, and 937 cm^-1^. These peaks match the literature values for anhydrous biogenic guanine(30). As described in *Flabellina iodenea*(5), guanine’s high refractive index in the [100] direction (∼1.83) allows high intensity reflections to be achieved with only a few layers. While the presence of guanine in other species of nudibranch is not unexpected, we believe that the diversity of colors that it is able to produce is remarkable, even within a single individual. Interestingly, for the two *Hypselodoris* species, guanine was associated with a pigment peak (Raman shift of 1155 cm^-1^) which fell over time under laser illumination, relative to the guanine peaks (see **Supplementary Figure 4**), whilst for the two aeolid species, the presence of guanine crystals is associated with an unidentified fluorescent compound (see **Supplementary Figure 5**). These additional components verify that structural color in these species works in synergy with pigmentary coloration. Based on our Raman characterization, the color mechanism is the same in these two groups; however, the question still remains as to whether the evolution of structural color in these groups emerged in a common ancestor or via convergent evolution. Due to the lack of a fossil record, it is challenging to infer divergence dates between nudibranch groups. Layton et al. calibrated a mitochondrial COI rate for chromodorid nudibranchs of up to 4.9% per million years per lineage, using the Isthmus of Panama closure (∼2.8 Ma)(31), whilst Carmona et al. report COI divergences of up to 16% between aeolid genera(32). However, such divergences cannot be directly extrapolated to deeper nodes, such as the dorid-aeloid split, due to rate variation over deep time. Jörger et al. place major Euthyneura splits—including opisthobranch diversification—in the Mesozoic (Triassic– Jurassic)(33). Together, these studies imply that the dorid–aeolid nudibranch split may have occurred during the Mesozoic, probably Jurassic to Early Cretaceous, though targeted molecular clock analyses are needed.

### Guanine multilayer stacks in *C. annae*

In order to characterize the nanoarchitecture responsible for the structural coloration, cryo Plasma-FIB SDB was conducted on a region of blue *Chromodoris annae* dorsal mantle tissue. Two regions of interest were imaged (**Figure 3, Supplementary Figure 6**), finding stacks of electron-dense material in both cases, lying from 2 *μ*m up to 20 *μ*m from the surface of the sample. Correlating this with the Raman spectra shown in **Figure 2**, we conclude that these crystals are biogenic guanine. In 2D, these electron-dense regions appear as rods (**Figure 3A**); however, when reconstructed in 3D (**Figures 3B, Supplementary Figure 6**), the electron-dense regions have a clear plate-like morphology (as expected for guanine crystals). The in-plane shape is faceted and anisotropic, forming a mixture of elongated hexagons and rhombi with lengths 0.1 – 1.5 *μ*m and widths of 0.1 - 0.5 *μ*m. These platelets form stacks of up to 16 crystals (with an average of 6 ± 3 platelets per stack), with a less electron-dense material separating them. The stacks are enclosed within inner membranes (**Figure 3A** red), which are in close contact with each other within an outer membrane or envelope (**Figure 3A** yellow). The diameter of the inner membrane-enclosed volume bounding the stacks varies from 1 - 3 *μ*m, whilst the outer membrane encloses a volume which is ∼ 8 *μ*m in its long direction and 5 *μ*m in its short direction. The envelopes containing volumes of crystalline stacks are interconnected (**Supplementary Figure 6**), forming a network that extends over 10 *μ*m in all three dimensions. The mean thickness of a platelet is 53 ± 10 nm, ranging from 39 to 68 nm, whilst the mean spacing of the platelets is 70 ± 24 nm, giving a total periodicity of the multilayer stacks of ∼120 nm.

**Figure 3.**
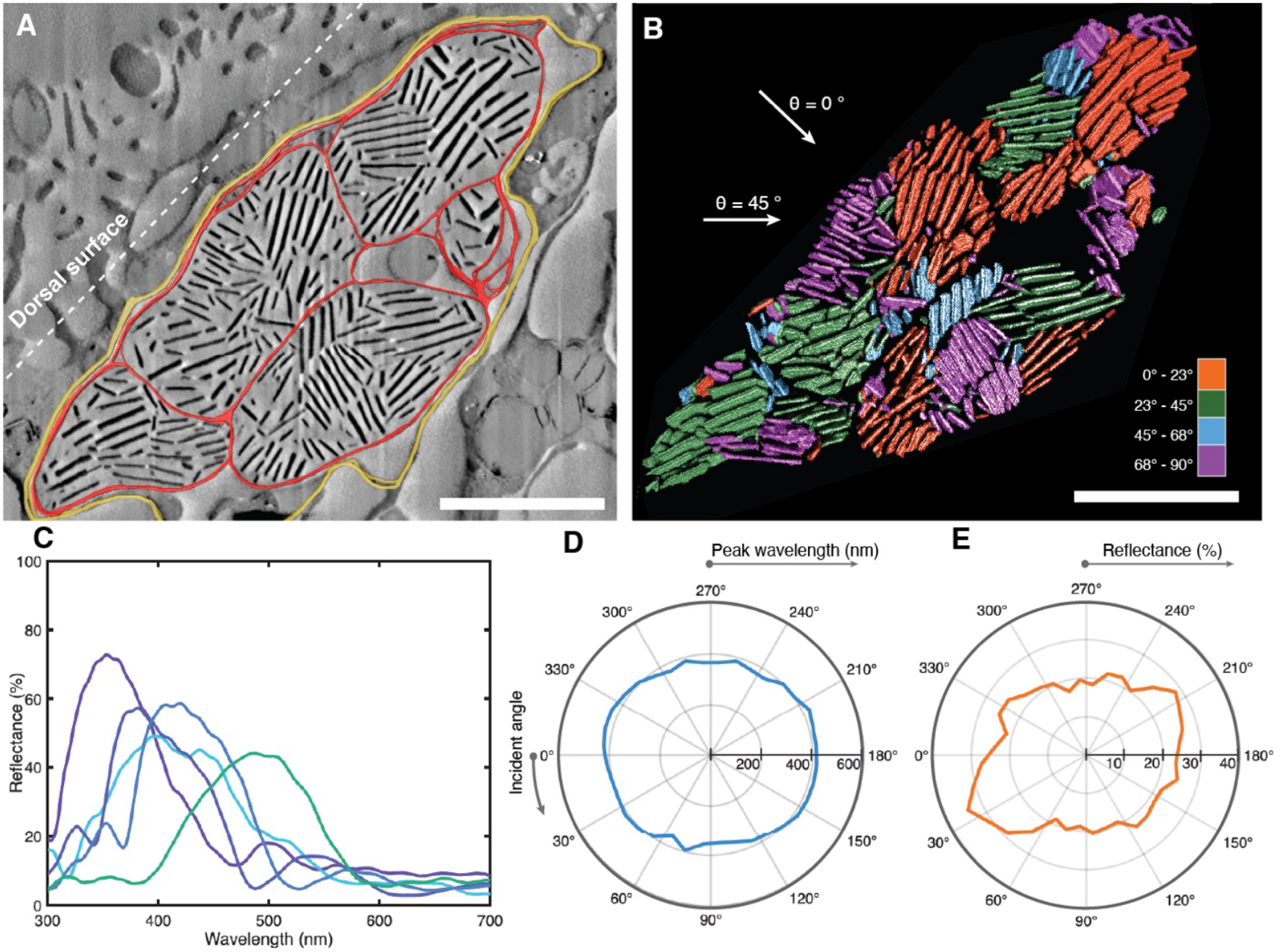
Nanoscale characterization of tissue from the blue dorsal region of *Chromodoris annae* with cryo-Plasma-FIB SDB and optical modelling. The volumes imaged here are located 2 to 20 *μ*m from the surface of the sample. A) Combined SEM image with a 3D reconstruction of outer membrane (yellow) and inner membranes (red) superimposed. Inner membranes contain multiple stacks of electron dense crystals with apparently random orientation ranging from 1 to 16 crystals thick. Scale bar, 3 *μ*m. B) 3D reconstruction of the electron dense crystals in A), showing the crystals are plate like in morphology. Stacks are colored by their orientation. Scale bar, 3 *μ*m. C) FDTD simulations of the reflectance spectra of individual guanine platelet stacks isolated from the volume in (B), taken at normal incidence with respect to the orientation of each stack. Spectra show sharp reflectance peaks (full width at half maximum ranging from 80 – 120 nm) with peak positions ranging from 350 – 500 nm and reflectance of up to 73%. D) Polar plot of reflectance spectra peak position vs incident angle θ when illuminated over the entire reconstructed volume in (B). The incident beam is rotated around the volume (angles are indicated in (B)), demonstrating that spectral peak position is relatively angle independent – remaining between 350 nm and 430 nm at all angles of incidence. E) Polar plot of reflectance spectra maximum amplitude vs incident angle when illuminated over the entire reconstructed volume in B, corresponding to the peak positions in d. The amplitude varies from 17 – 34 %.

The optical response of a single stack of guanine platelets, such as those found through FIB SEM imaging here, can be approximated to that of a standard multilayer reflector. When the plate orientation, thickness, and spacing are uniform within the structure, reflection intensities up to 100% can be achieved with only a few layers due to the high refractive index contrast of the platelets(12). For this reason, the multilayer is an efficient structure to generate bright and brilliant colors. To verify that the structure imaged in **Figure 3B** is responsible for the observed optical properties, we simulated the optical response of individual stacks within the volume by performing finite-difference time-domain (FDTD) simulations on the volume reconstructed from FIB SEM imaging (Methods). The simulated spectra contain well-defined, bright (up to 73%) reflectance peaks ranging from 350 – 500 nm (**Figure 3C**), closely matching the experimental spectra shown in **Figure 2L**. Our simulations confirm that this structure is responsible for the blue coloration seen *in vivo*.

A homogeneous multilayer structure gives rise to an angle-dependent response, where a blue shift occurs as the angle of illumination is increased. The wavelength at which the reflection is maximized in a multilayer can be calculated simply using Bragg’s law (λ=2ndcosθ) where n = refractive index contrast between the layers, d = lattice constant, and θ = angle of incidence(34). Consequently, the color reflected from a perfect infinitely large individual stack shifts to lower wavelengths at higher incident angles (**Supplementary Figure 7B**). Here, however, we observe that *Chromodoris annae* maintains its bright blue appearance (**Figure 1D)** when viewed from any angle by varying the orientation of the multilayer stacks in space. In the reconstructed volume in **Figure 3B**, as an example, the differently oriented multilayer stacks (shown in different colors) are broadly and evenly distributed between 0° and 90° relative to the animal’s surface, with an average angle of 50° ± 30° measured across 25 stacks (see **Supplementary Figure 8** for the full platelet angle distribution). The misorientation of the crystal stacks means that, whatever the viewing angle, there will be a multilayer stack which is oriented perpendicular to the incident light, and constructive interference will thus achieve blue reflectance. To confirm this effect, we simulated the optical response of the entire volume in **Figure 3B**, whilst varying the incident angle θ, finding that at all values of θ, the peak wavelength of reflected light lies between 350 and 430 nm (**Figure 3D**). The reflected intensity in this configuration varies from 17 - 34% (**Figure 3E**). Variation in stack orientation to homogenize color from multilayer stacks has been reported in other systems, such as the blue ringed octopus(35). Such a hierarchical structure achieves angle-independent coloration from an intrinsically angle-dependent photonic structure.

We have also shown through 2D block-face SEM imaging that the dorsal surface of the dorid nudibranch *Hypselodoris bullockii* contains similar misoriented guanine crystal stacks (2 - 10 platelets per stack), confined within inner and outer envelopes (**Supplementary Figure 9A, B**). In this case, the mean platelet thickness is 49 ± 7 nm and the mean spacing between platelets is 69 ± 20 nm giving a total pitch of 118 nm. This relatively smaller pitch shifts the constructive interference condition to shorter wavelengths, explaining why the reflectance peaks in **Figure 2K** are blue-shifted from those of *C. annae* in **Figure 2L**. Additionally, imaging the cerata of the aeolid nudibranch *Berghia stephanieae* (**Supplementary Figure 9C-F**) in this way demonstrated an array of guanine platelets with a mean platelet thickness of 150 ± 30 nm. Here the platelets are stacked around the border of envelopes (∼10 *μ*m from the surface of the ceras, around the entire perimeter). The space between these platelets (which is far smaller than in the dorid species, relative to plate thickness), and their orientations are not well defined. This lack of structural coherence explains the large variation in coloration and textured appearance of the domains in this species (**Figure 2H, P**), resulting in an overall white appearance.

To give an idea of how small changes in the guanine platelet thickness and spacing can give rise to different optical responses across the visible range, we use a transfer matrix method(36) to compute guanine multilayer stacks with varying plate thickness and spacing. **Supplementary Figure 7C** shows that by varying plate thickness from 50 to 90 nm and spacing from 50 to 110 nm for multilayer stacks containing 6 platelets, reflectance peaks from across the entire visible wavelength range can be achieved. These simulations support our hypothesis that the nudibranch species studied here generate a wide range of colors by varying the thickness and stacking pitch of guanine platelet stacks. On a larger scale, nudibranchs utilize these structurally tuned stacks to give pixels (domains) of different color, which combine to give a range of colors based on their statistical distribution(37). In other words, ultrastructural tuning determines the available color palette on a microscopic scale, whilst the proportion of pixels with each ultrastructure determines the macroscopic color. Their angular distribution permits matt rather than iridescent coloration(37).

Although the range of structural colors described here is wide, it should be noted that in many species these colors act in combination with pigmentary color. *Flabellina iodinea* is described as having ‘silvery reflectance’ which enhances pigmentary color(5). However, silvery reflectance should only enhance pigmentary coloration if the silvery reflector lies below the pigment (such that light has multiple opportunities to be absorbed). Here, the photonic structure in **Figure 3A, B** was located at the surface of the animal and therefore lies above any pigments present in the body. Therefore, we suggest that guanine crystal stacks are responsible for the generation of specific hues in the species studied here. So, although these structural colors act synergistically with the pigmentary coloration, they not only enhance colors but also are responsible for generating them.

## Conclusions

Through optical and ultrastructural imaging, we have shown that nudibranchs from across the dorid and aeolid groups utilize guanine crystals for highly variable structural coloration. Based on imaging and simulations, we suggest that variation in color of granules comes from a variation in guanine platelet size and stacking pitch. Different species will utilize different length scales of guanine platelets to generate different colors. For example, *H. bullockii* exhibits structurally colored granules in the near-UV range, due to a smaller pitch of guanine platelet stacks, whilst the white region of *B. stephanieae* displays granules across the visible range up to the red region, because of a greater pitch. On top of this multilayer pitch modulation, we propose that the nudibranchs *C. willani* and *C. annae* have developed a novel method for producing brilliant white, which utilizes an array of guanine multilayer stacks in combination with a scattering material. The misalignment of stacks in *C. annae* allows for angle-independent coloration, which may be key for the aposematic function of color in these slow-moving animals, allowing them to deter predators from any viewing position.

Behavioral studies would be required to understand the aposematic role of guanine here, but it certainly imparts very bright colors onto the skin of these animals (up to 80% reflectance for *C. annae*) in a variety of striking colors. Subtle structural changes in multilayer guanine reflectors— capable of generating a broad spectrum of colors — may have played a key role in the evolution of color diversity among nudibranchs in the two clades described here. Such architectures, not only allow for bright colors to be achieved, but also for multiple colors in the same species made from the same material, increasing the possibility of pattern diversity.

## Methods

### Phylogenetic tree

A phylogenetic tree was constructed using phyloT (https://phylot.biobyte.de), a web-based tool that generates trees based on the hierarchical taxonomy in the NCBI Taxonomy database. A list of the 14 relevant species and their associated NCBI Taxonomy IDs was used as the input. phyloT retrieved the taxonomic lineages for each entry and constructed the tree using these lineages and their hierarchical relationships.

NCBI taxonomic information integrates information from a variety of biological data sources, including genetic studies, morphological traits, databases and published literature. Importantly, phyloT does not use sequence data or apply clustering algorithms. Taxa are instead grouped and ordered based on their positions within the taxonomy database.

The tree was exported in Newick format for visualization using iTOL (Interactive Tree of Life) (https://itol.embl.de) and subsequently reproduced using Adobe Illustrator.

### Photography and digital microscopy

A Keyence VHX-7100 digital microscope was used to take high magnification images of live sea slugs (see **Supplementary Figure 10** for details). A Nikon D7500 SLR camera was used to take macroscopic images of the sea slugs.

### Optical imaging and spectroscopy

Micro-spectroscopy was carried out using a customized Zeiss Axio Scope A1 microscope, fitted with a Pixelink PL-D725CU-T high frame rate camera, which was color calibrated with a white diffuser (Labsphere USRS-99-010). A Zeiss HAL100 halogen lamp was used as a light source. For acquiring spectra, the microscope was coupled to an Avantes AvaSpecHS2048 spectrometer using an Avantes FC-UVIR50 (50 *μ*m core diameter) multimode optical fibre. Spectra were taken using water immersion objective lens 40x (Zeiss, WN-Achroplan 40x (NA 0.75, FWD 2.1 mm)). Spectra were referenced to a protected silver mirror (PF10-03-P01, with >97.5% reflectance for 450nm - 2*μ*m). Granule diameters were measured as the mean of 35 discrete granules (optically distinct islands of a single color).

The samples were placed on a glass slide and a coverslip was placed on top. Samples were then imaged on a Keyence VHX-7100 digital microscope before being transferred for spectra. Seawater was used as the medium between the water immersion objective and cover slip to ensure index matching. Spectra were taken in the Koehler illumination configuration, with the aperture diaphragm close to fully closed to reduce the collection of stray light from undesired areas. An image of the fiber collection area was taken prior to spectra using an external lamp, and a circle representing this area overlayed to on the PixelLink imaging software to accurately know from which area spectra were being taken (Supplementary **Figure 11)**.

### Raman spectroscopy

Raman spectroscopy was performed on cryo-cut, unfixed, unstained room temperature 14 *μ*m thick sections of sea slug tissue. Samples were cut at -20 ° c using a Leica CM3050 S cryostat (aside from *H. bullockii* and *B. stephanieae* in which case samples were cut at room temperature). An alpha300 R – Raman Imaging Microscope (Oxford Instruments) equipped with a UHTS spectrometer was used to take Raman spectra. Spectra were taken with a 100× objective (Nikon) in confocal mode. A near-IR laser (*λ* = 785 nm) was used for all species due to fluorescence induced by the green laser in aeolid samples. Green laser (532 nm) Raman spectra are shown in **Supplementary Figure 12**. For each species the desired region of interest was located on the sections using a Keyence microscope in full ring mode (see SI). The sample was then transferred to the Raman microscope and the desired region of interest was measured in at least 3 positions with an integration time of 60 seconds for each. These positions were averaged and processed using the Project *FIVE*+ software (Oxford Instruments). For Raman mapping an integration time of 5s was used due to time constraints.

### FIBSEM

This data was taken on a Thermo Fisher Hydra Bio Plasma-FIB. 3D reconstruction was done using both the Thermo Scientific Amira Software and ORS Dragonfly software. The sample is a region of blue dorsal tissue of *Chromodoris annae* high pressure frozen in hexadecane as a cryoprotectant. The desired areas of tissue were cut from the body of a sedated slug with a scalpel and under a stereo microscope (Zeiss). Immediately after dissection the sample was placed into a B gold-coated copper freezer hat (BALTIC preparation, Wetter, Germany) and 10 wt% dextran (Sigma, 31390) was added as a cryoprotectant. A second freezer hat was added in a mirror configuration to enclose the sample, giving a total cavity thickness of 0.6 mm. Within approximately 5 minutes of decollation the sample in this sandwich combination was cryo-immobilized using a high-pressure freezer (HPM100, Leica Microsystems).

### FIBSEM data acquisition

Sequential Plasma Focused Ion Beam (PFIB) milling and imaging (also often referred to as ‘Slice & View’) were conducted using the Helios 5 Hydra DualBeam system (Thermo Fisher Scientific, Inc., Waltham, MA). Secondary electron images were acquired with the Thru-lens detector (TLD) in immersion (UHR ultra-high resolution) mode. Samples mounted on a 35° pre-tilt shuttle were loaded onto the microscope cryo stage. Experiments utilized Argon ions at 30 kV. A protective, non-conductive platinum-organic coating was deposited on the sample surface using a Gas Injection System (GIS), sandwiched between two sputter-coated conductive platinum (Pt) layers before milling trenches. Sputter coating was performed at 12 keV and 250 nA, with layer thicknesses of approximately 5-10 nm, while the GIS coating was around 1 *μ*m thick.

Upon identifying an area of interest, a trench was milled using 30 kV, ∼7 nA, followed by further polishing with 2 nA and 0.74 nA currents. This process enabled rapid verification of the area of interest within the milled region and facilitated quality assessment. Subsequently, a trench was milled on the left side of the initial front trench to mitigate charging effects, which typically manifest as “dark flares” from the left side of the image. A portion of the dark region near the left trench was intentionally included in the imaging area to serve this purpose.

Volume data acquisition was performed using Thermo Scientific™ Auto Slice and View 5 software under the following conditions: Milling parameter – 30 kV, 200 pA; slice thickness – 20 nm; SEM landing energy – 1.2 keV; beam current – 13 pA; dwell time – 50 ns; line integration – 150; lateral pixel size – 5 nm × 5 nm.

### FIBSEM Data processing

The data was imported into Thermo Scientific™ Amira software for post-processing and aligned using a rigid translational approach. Denoising was initially performed using a simple Gaussian filter with settings of 1,1 and a kernel size of 2. To further enhance the separation of crystals from the surrounding tissue, a Non-Local Means denoising filter was applied with the following settings: Search Window size = 30 px, Local Neighborhood = 2 px, and Similarity Value = 0.4. The Match Contrast module was utilized to normalize contrast brightness across the entire stack.

Segmentation of the crystals was conducted in Amira’s classic segmentation room and ORS Dragonfly. In Amira this was done using a combination of the Threshold module and manual refinement of the resulting selection. For instances where multiple color volumes were displayed, a labeling step was performed to assign different colors to objects without common pixels. The Label Analysis module was used to extract volume and surface measurements of the crystals. In Dragonfly segmentation was conducted using a custom uNet model, this model was trained using 3 SEM images with thresholding and subsequently manually corrected.

Finally, Volume Rendering was applied to the segmented data to visualize it in 3D, and all measurements were performed in 3D using Amira’s 3D measuring tool. Both stacks, containing 13 and 295 slices respectively, were processed using the same protocol described above.

### Freeze substitution

The freshly cut tissue (cerata of *B. stephanieae*, pink skirts of *H. bullockii*) of nudibranchs were sandwiched individually between two type B gold coated copper high pressure freexer carriers (BALTIC preparation, Wetter, Germany) with the addition of sea water. The carriers were positioned in a mirror combination to allow a total cavity thickness of 0.6 mm. The sandwiched samples were cryo-immobilized in an EM ICE high-pressure freezing machine (Leica Microsystems, Vienna, Austria) within 10 minutes after dissection. Freeze substitution was carried out in an automatic freeze substitution unit (Leica Microsystems, AFS). The samples were firt transferred under liquid nitrogen to tubes containing freeze substitution medium composed of 2 wt% OsO4, 0.5 wt% glutaraldehyde, and 1.5 wt% water in acetone. Tubes were placed in AFS which was set at −120 °C. They were subsequently warmed to −85 °C at a rate of 17.5 °C/h and held at −85 °C for 103 h, then warmed to −20 °C at a rate of 7.2 °C/h, followed by a hold at −20 °C for 12 h, and a final heating to 4 °C at a rate of 2.7 °C/h followed by holding at 4 °C for 24 h.

The samples were rinsed with acetone, infiltrated with Embed 812 epoxy resin for 3 days. The resin polymerization was done for 24 hours at a curing temperature of 65 °C.

### Finite Difference Time Domain (FDTD) simulations

FDTD simulations were performed with a commercial solver (Ansys Lumerical). The 3D cryo-FIB-SEM data was imported as an isotropic material and embedded in a 2D FDTD simulation with PML boundary conditions in all directions, a mesh size of 15 nm, and a total simulation time of 2 ps. The imported structure was illuminated with a broadband diffracting plane wave and the power was monitored with a linear DFT monitor placed behind the source in the incident direction to collect reflected light (see **Supplementary Figure 13**). For simulating the reflectance of individual guanine domains, source and detector were positioned as close as possible to the structure to simulate high-NA collection and the simulation window was 2 *μ*m × 2 *μ*m. To calculate the angular dependence of the whole structure, the source and detector were placed approximately 5 *μ*m away from the centre of the structure, which is rotated by a total of 360º in steps of 10º, in a simulation area of 11 *μ*m × 11 *μ*m. At each angle of rotation, the 2D simulation was run over all z-planes of the image dataset and the resulting spectra were averaged and smoothed with a savitzky-golay filter. The maximum reflectance of each resulting spectrum was retrieved, and the centre wavelength was calculated by fitting a gaussian to the averaged and smoothed reflectance spectrum. Refractive indices were n = 1.83 for the guanine layers and n = 1.34 for the surrounding tissue.

### Transfer matrix simulations

The reflectance spectra used to generate **Supplementary Figure 7** were obtained via numerical simulations using a modified StackModel routine from the open-source PyLlama Python library(36). In **Supplementary Figure 7b**, simulations were performed by sweeping the incidence angle from 0° to 45° for a multilayer system consisting of N = 10 stacks, each composed of platelets with a thickness of 55 nm and a spacing of 70 nm. In **Supplementary Figure 7c**, reflectance was simulated across the visible-UV spectral range, using a leptokurtic distribution of incidence angles with the center of mass around normal incidence. The model assumed an isotropic multilayer system, also consisting of N = 10 stacks, where each platelet had a thickness varying between 50 nm and 90 nm, separated by gaps ranging from 50 nm to 110 nm. To better mimic experimental conditions, Gaussian white noise was added to the platelet thickness, with a threshold of less than 10%. In all cases, the system followed a sandwich heterostructure composed of two materials with refractive indices n_A = 1.83 and n_B = 1.34 (background), respectively.

### Handling and ethics

1-2 individuals of each species were studied. All species were bought from online aquarium shops (Rifix, Germany, Coral & Fish Store and De Jong Marinelife, Netherlands) apart from *S. neapolitana* which was collected in an intertidal zone in the Eastern Atlantic. Dorid nudibranchs originate from the western Indo-Pacific, and this study has been acknowledged by the Republic of the Philippines Department of Biodiversity and Natural Resources, Biodiversity Management Bureau. Animals were studied immediately after arrival. Tropical species were kept at 26 ºC and temperate species were kept at 18 ºC in separate aquaria. All specimens were treated with best practices; this includes sedating of animals prior to experiments and regular replacement and aeration of seawater. None of the listed species are protected under CITES, IUCN, or national endangered species acts.

## Acknowledgments

This work was supported by the ERC BiTe ERC‐2020‐CoS-101001637, EU Horizon 2020 program (H2020-MSCA-ITN-2019) grant N 860125 “BEEP”, The Marie Curie Foundation FucAd, Grant agreement ID: 101155041. We would also like to acknowledge Thermo-Fisher for the electron microscopy undertaken at their facilities. S.P.P. thanks Dr. Kevin Vynck the hospitality and computational support for the numerical simulations (SALTO program). We also acknowledge Dr Yu Ogawa for his assistance with sample preparation and imaging.

