## Supplementary information for "Nudibranch color diversity shares a common origin in guanine photonic structures"

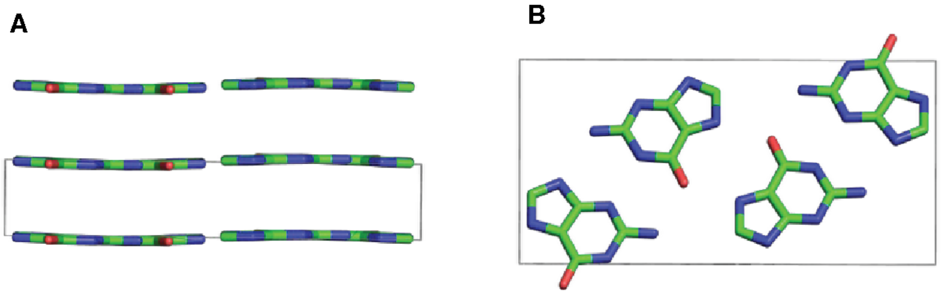


**Supplementary Figure 1**

Monoclinic β polymorph of guanine showing (A) the view along the b-axis (edge-on) of the stacked molecular layers and in (B) a top view of the bc molecular plane. Blue = N, Green = C, Red = O.

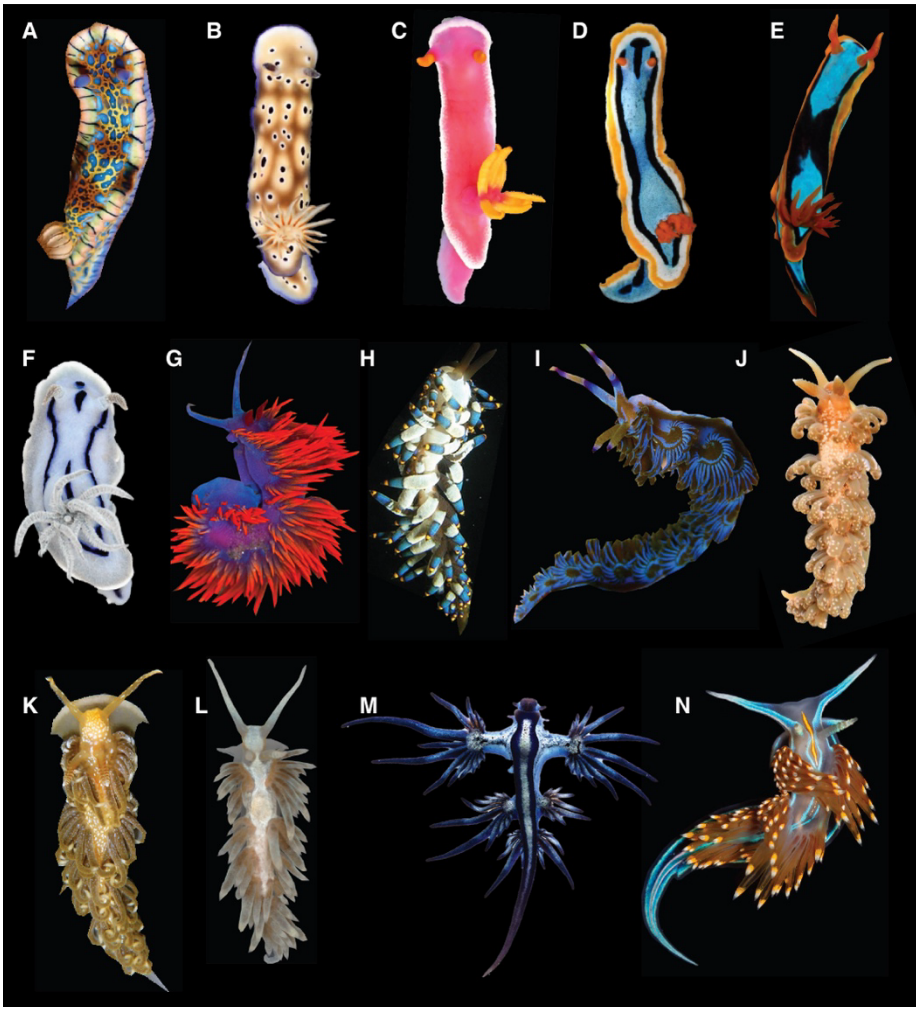


**Supplementary Figure 2**

Photographs of the dorid and aeolid nudibranch species included in **Figure 1**,

A) *Hypselodoris acriba*, Dan Schofield. Reproduced from iNaturalist, under CC BY 4.0, http://creativecommons.org/licenses/by/4.0/
B) *Hypselodoris tryoni*, Samuel Humphrey
C) *Hypselodoris bullockii*, Samuel Humphrey
D) *Chromodoris annae*, Samuel Humphrey
E) *Chromodoris westraliensis*, Zoe Richards et al. Reproduced from Wikimedia commons., under CC BY 4.0, http://creativecommons.org/licenses/by/4.0/
F) *Chromodoris willani*, Samuel Humphrey
G) *Flabellina iodinea*, Jerry Kirkhart. Reproduced from Wikimedia Commons, under CC BY 2.0, https://creativecommons.org/licenses/by/2.0/
H) *Trinchesia yamasui,* Chika Watanabe. Reproduced from Wikimedia Commons, under CC BY 2.0, https://creativecommons.org/licenses/by/2.0/
I) *Pteraeolidia ianthina,* Sylke Rohrlach. Reproduced from Wikimedia Commons, under CC BY 2.0, https://creativecommons.org/licenses/by/2.0/
J) *Spurillla neapolitana*, Samuel Humphrey.
K) *Spurilla braziliana,* Robin Gwen Agarwal. Reproduced from iNaturalist, under CC BY 4.0, https://creativecommons.org/licenses/by-nc/4.0/
L) *Berghia stephanineaei*, Xianglian He.
M) *Glaucus atlanticus*, Taro Taylor. Reproduced from Wikimedia Commons, under CC BY 2.0, https://creativecommons.org/licenses/by/2.0/
N) *Hermissenda opalescens*, Nathan Foster. Reproduced from iNaturalist, under CC BY 4.0, https://creativecommons.org/licenses/by/4.0/

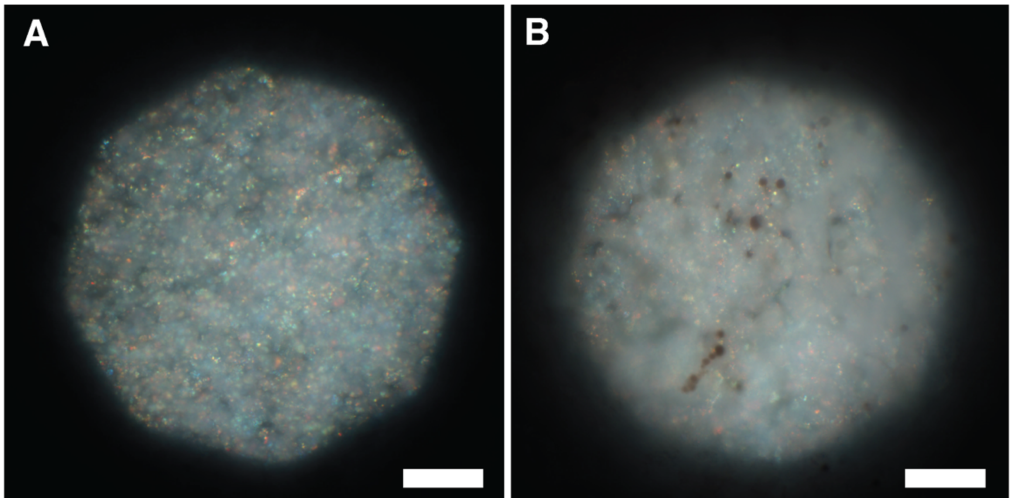


**Supplementary Figure 3**

Upright optical microscope (Zeiss Axioscope, 40x WI objective) of an area of white tissue of *Chromodoris willani.* In (A) the image is recorded, then the pixellated area is in focus, showing small, colored grains across the visible spectrum. (B) shows the same area as (A) but focusing above the pixelated region, showing a cloudy white color and small pigment spots. Both Scale bars are 30 $\mu$m**.**

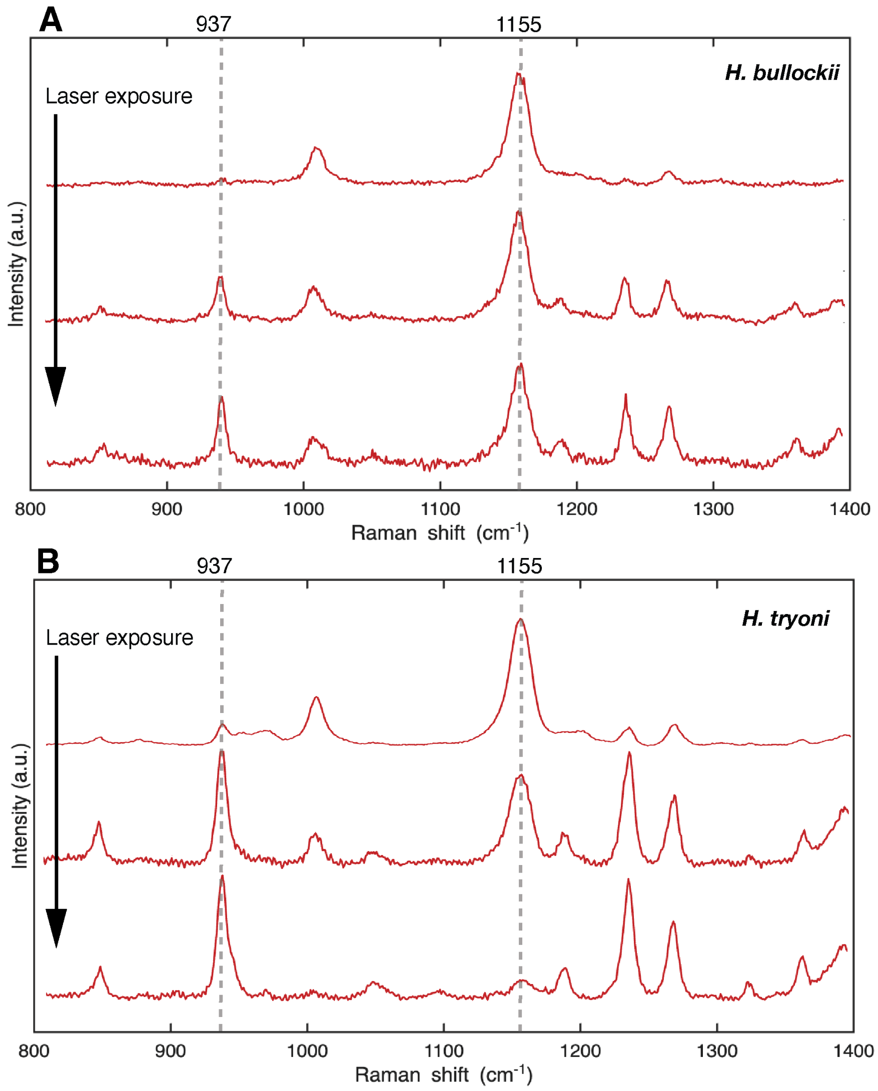


**Supplementary Figure 4**

Raman spectra of iridescent regions located close to the dorsal surface in histological sections of the 2 *Hypselodoris* species studied, specifically in (A) *H. Bullockii* and in (B) *H. tryoni.* The two peak positions highlighted show one of the peaks for biogenic guanine (937 $cm^{-1}$), and a second peak (1155 $cm^{-1}$), which we attribute to an unidentified pigment. The relative heights of these peaks shift in favor of the guanine peak with increased laser exposure, likely due to bleaching of the pigment.

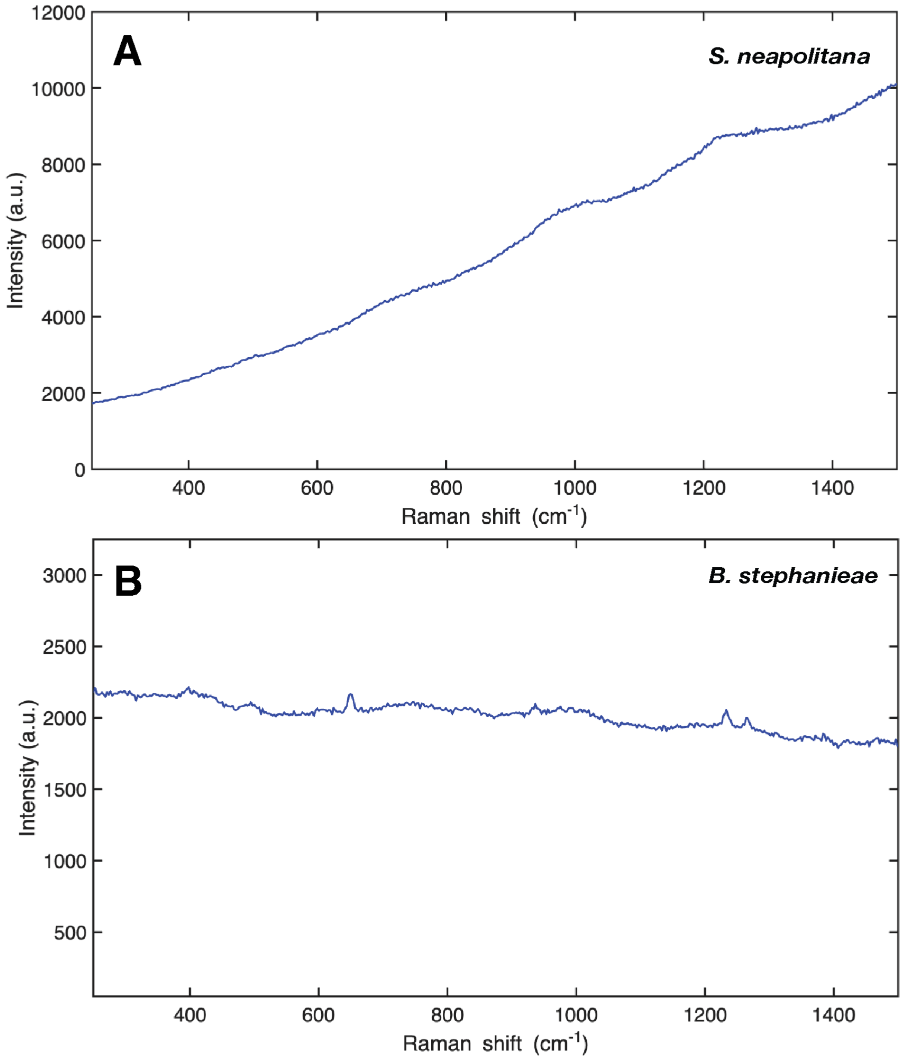


**Supplementary Figure 5**

Raman spectra of iridescent regions located close to the dorsal surface in histological sections of the 2 aeolid species: (A) *S. Neapolitana* and (B) *B. stephanieae* studied. Biogenic guanine peak positions are masked by induced fluorescence. These spectra were taken with a green laser (532 nm).

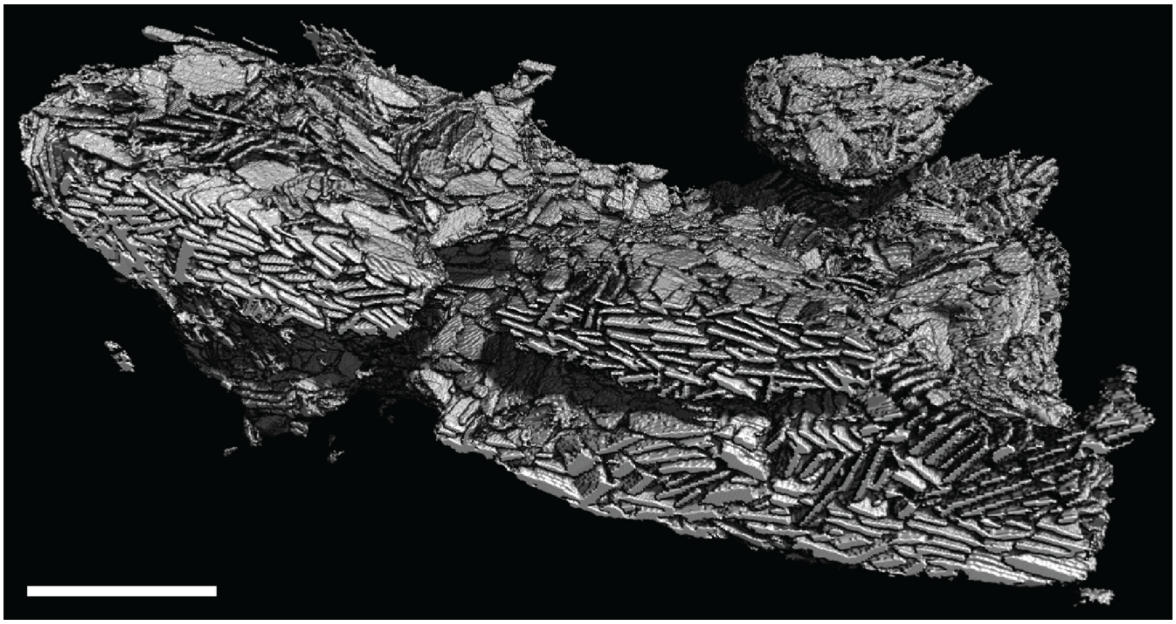

**Supplementary Figure 6**

3D reconstruction of a region of guanine multilayer stacks on a region of blue *Chromodoris annae* dorsal mantle tissue, showing the crystals are plate-like in morphology. Stacks typically contain 2 - 5 crystals, and the crystal-containing regions extend in a connected network which is greater than 10 $\mu$m in all directions. Scale bar, 2 $\mu$m.

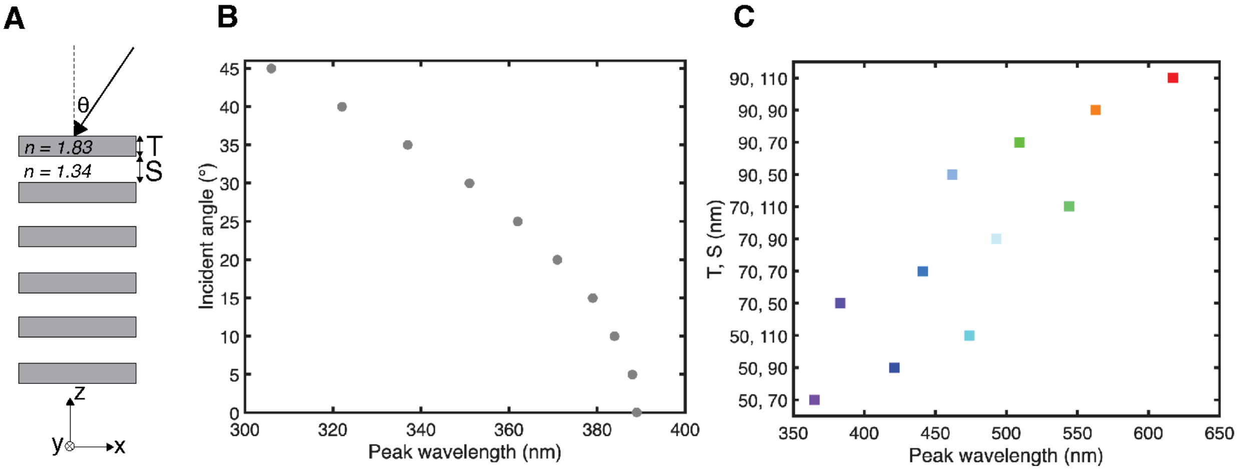


**Supplementary Figure 7**

2D simulations conducted using the transfer matrix method (PyLlama) for a multilayer stack. (A) Schematic showing key parameters used for 2D simulated spectra. These are: platelet thickness (T), platelet spacing (S), incident angle (θ) and refractive indexes of 1.83 and 1.34 respectively.

B) Plot of spectral peak wavelength vs incident angle of simulated reflection spectra for guanine platelet stacks with parameters taken directly from **Figure 3** (T = 55 nm, S = 70 nm, N = 6), incident angle is varied from 0 – 45º showing a blue shift of the reflectance peak into the UV range with increasing incident angle.

C) Plot of spectral peak wavelength with varied plate thickness T and spacing S for simulated reflection spectra from a multilayer stack of 6 layers of material. plate thickness T varies from 50 – 90 nm and plate spacing S varies from 50 – 110 nm. This simulation demonstrates that peaks across the visible range with reflectance values of over 80% are possible from just five layers of material, and that by varying the dimensions of these structures any reflected color can be achieved. This simulation was conducted using PyLlama software (see methods section).

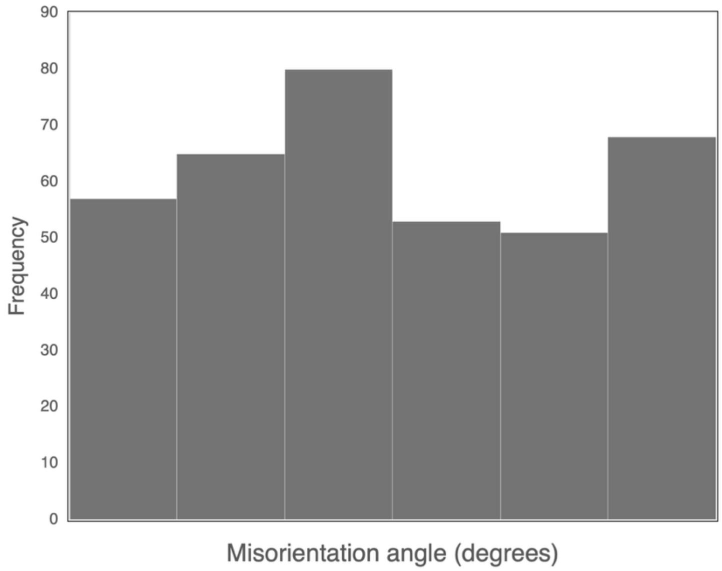


**Supplementary figure 8**

Histogram of misorientation angles of guanine nano-platelets contained in the volume in main text **Figure 3**, relative to an arbitrary plane XY.

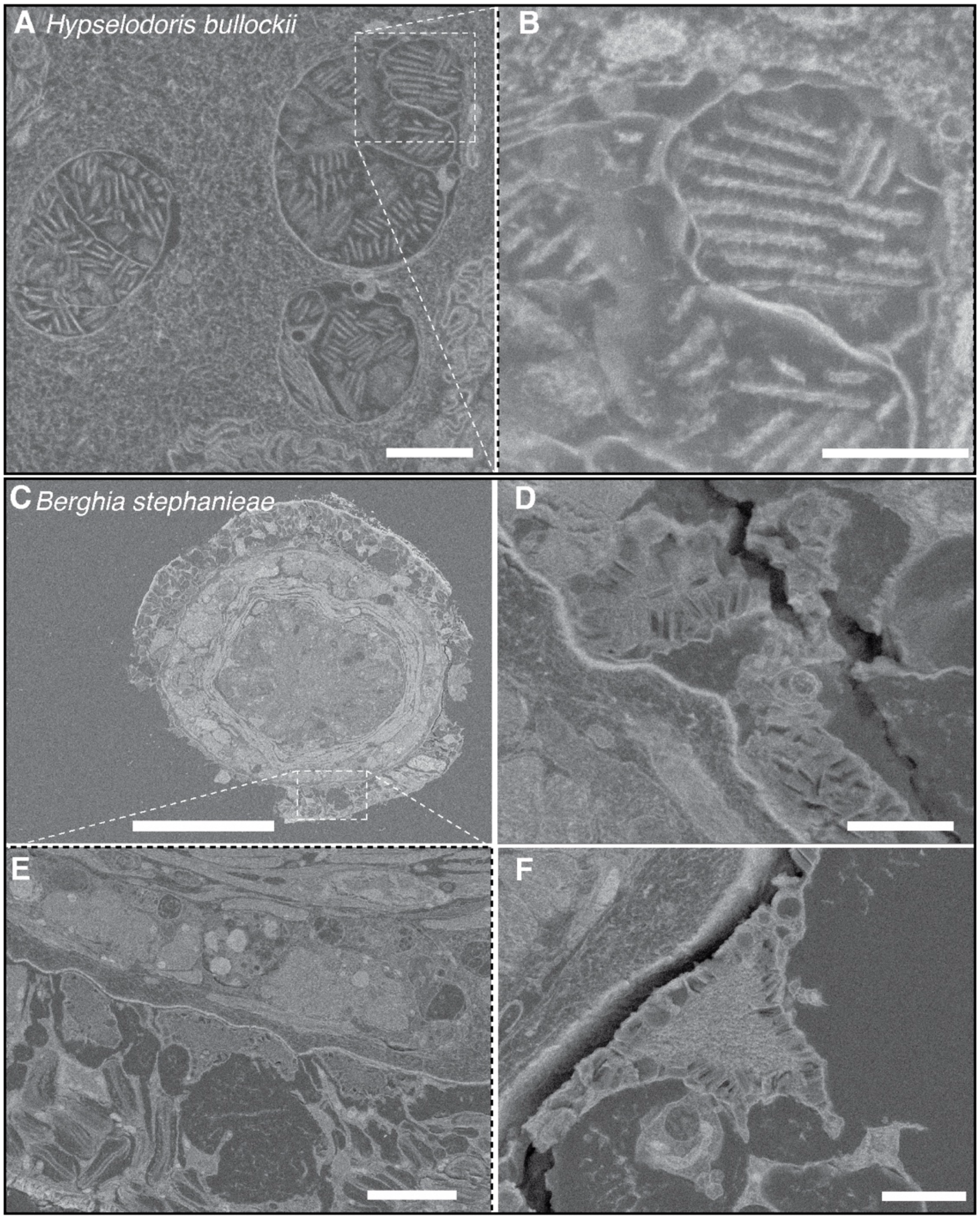


**Supplementary Figure 9**

Block-face SEM images of *Hypselodoris bullockii* (A, B) and *Berghia stephanieae* (C-F). In *H. bullockii* there are guanine platelet stacks containing 2-10 platelets per stack, contained within inner envelops, inside a larger envelope. In *B. stephanieae* platelets form stacks at the borders of membranes, which defines the orientation of the stacks. However, the orientation within stacks is more random as compared to *H. bullockii.* Guanine platelets can be found around the entire perimeter of the ceras, approximately 10 μm from the surface. Scale bars: 2 μm, 1 μm, 50 μm, 2 μm, 5 μm, 2 μm.

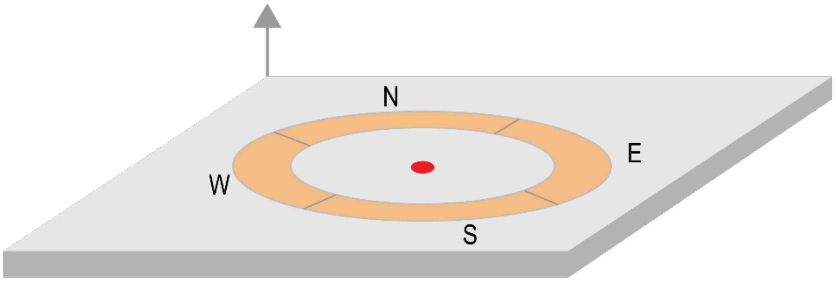


**Supplementary Figure 10**

Schematic showing the illumination configuration of digital microscopy using a Keyence VHX-7100. The sample is illuminated either by the entire ring (full ring mode), or any one of the NESW directions (partial ring mode). The full ring mode is equivalent to dark field, in which the sample is illuminated from a ring of light on the outside of the objective, and light is collected from the center (marked by a red circle). This ring can be split into 4 separate illumination angles for 'partial ring' imaging

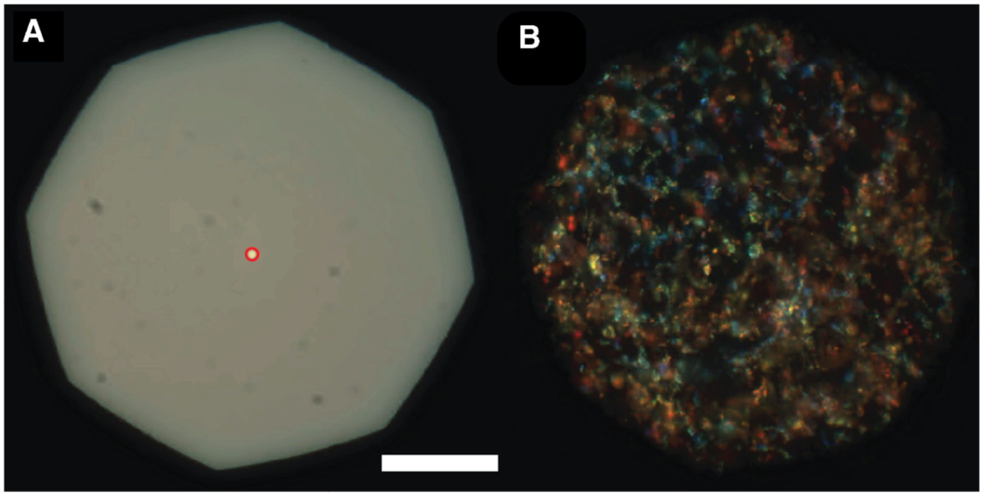


**Supplementary Figure 11**

Microspectroscopy setup. (A) A reference sample is illuminated with a lamp coupled into the optical fiber from the collection side to define the spectral collection area (bright spot). A red annotation is added on the PixelLink software to store the position. (B) Brightfield images of the sample allow correlating which region of the sample spectra are being collected from. Images taken on Zeiss Axio Scope A1 with a Zeiss W N-Achroplan 40x objective, using a 50 $\mu$m fiber. Scale bar, 30 $\mu$m.

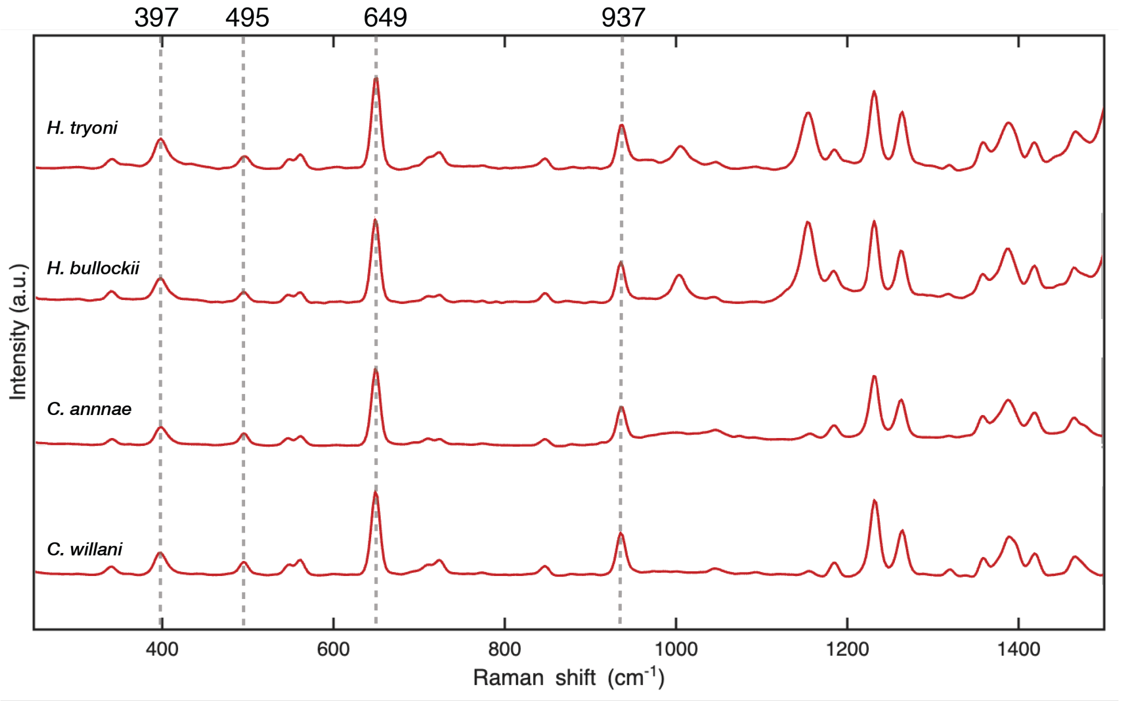


**Supplementary Figure 12**

Raman spectra of iridescent regions located close to the dorsal surface in histological sections of the 4 dorid species studied here. Highlighted peak positions correspond to literature values for anhydrous biogenic guanine. These spectra were taken with a green laser (532 nm).

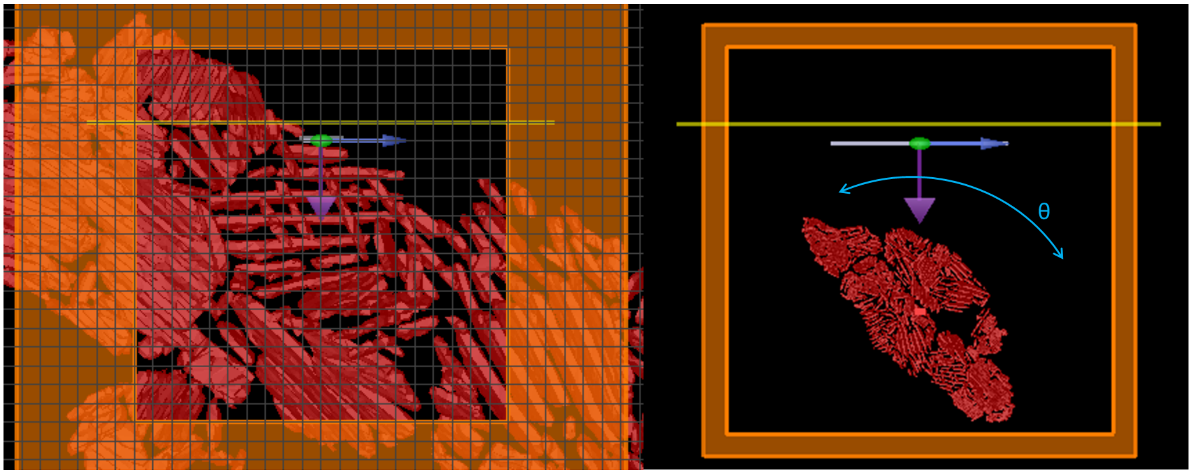


**Supplementary Figure 13**

Screenshot of the FDTD simulation setup in the x-y-plane. A plane wave source (green spot, travelling in the direction of purple arrow) is incident on the structure reconstructed from cryo-FIB-SEM data (red). The reflected power is monitored with a linear detector (yellow line). The 2D FDTD simulation with PML boundary conditions (orange) is centered around the structure. We simulate individual domains (left) or the entire structure (right), which is rotated by an angle θ, increasing counter-clockwise (shown here: θ = 225º).
